# Superposition eyes of diurnal and nocturnal hawkmoths are imperfect spheres and lack acute zones

**DOI:** 10.64898/2026.04.28.721382

**Authors:** Yash Sondhi, Ruchao Qian, John Paul Currea, Isabella Koushiar, Jacqueline Degen, Deborah Glass, Edward Stanley, Simon Sponberg, Ian J. Kitching, Akito Y. Kawahara, Jamie C. Theobald

**Affiliations:** Case Western Reserve University, Department of Biology, Cleveland, Oh; Florida International University, Miami, FL, USA; UCLA, Los Angeles, CA, USA; McGuire Center for Lepidoptera and Biodiversity, Florida Museum of Natural History, University of Florida, Gainesville, FL 32611 USA; University of Sussex, Falmer, Brighton BN1 9RH, UK; Natural History Museum, London SW7 5BD, UK; University of Oldenburg, Oldenburg, Germany; Georgia Institute of Technology, Atlanta, GA, USA

## Abstract

Superposition compound eyes improve sensitivity by pooling light from multiple facets and are widespread among nocturnal insects, including moths and beetles. Optical theory predicts that superposition eyes must be nearly spherical to form coherent images and therefore lack highly pronounced acute zones with high spatial resolution. To examine this, we imaged eyes of six hawkmoths (Sphingidae) spanning diurnal and nocturnal activity with high-resolution microcomputed tomography. Our automated pipeline created detailed eye maps quantifying morphological parameters. We measured local eye curvature (radial distance), interommatidial angle (Δφ), facet size and crystalline cone skewness (tilt of cone axes relative to the local surface normal). All species show more curvature in the dorso-ventral plane with flattening in the antero-posterior plane. However, their eyes still retain near-spherical curvature globally, with diurnal species showing greater distortion. For facet parameters, spatial acuity is generally highest (lowest Δφ) near the eye center and decreases gradually toward the periphery. However, overall variation in spatial acuity is low and these eyes lack distinct acute zones. Facet size gradually changes from center to periphery, increasing in some species and decreasing in the others. Cone skewness is present in all six species (0°-10°), but in two diurnal species of hummingbird hawkmoths it increases markedly in the posterior region (15°-30°) possibly compensating for regional eye flatness. This paper provides foundational data of ommatidial and eye shape measurements and advances our assumptions about how superposition eyes function.

## Introduction

Hawkmoths or Sphingidae are a diverse group of robust moths with over 1700 species and varied life histories (Kawahara et al. 2009; Kawahara and Barber 2015; Kitching et al. 2018). Their evolutionary history includes multiple transitions between diurnal and nocturnal adult lifestyles within a single clade (Kawahara et al. 2009; Akiyama et al. 2022). Hawkmoths are also model organisms for flight and sensory biology, able to navigate through cluttered vegetation in quite dim light (Stöckl and Kelber 2019), and discriminate color at night (Kelber et al. 2002; van der Kooi et al. 2021). Their visual ecology has been studied in depth for a handful of nectar-feeding, hovering species from the Macroglossinae and Sphinginae subfamilies (Stöckl and Kelber 2019) but not in Smerinthinae, which are predominantly non-hovering and non-nectar feeding as adults (Kawahara et al. 2009; Kawahara and Barber 2015; Kitching et al. 2018). The visual systems of nectar feeding and hovering hawkmoths have likely been optimized for three-dimensional flight control and rapid forward flight in their preferred light environment (Stöckl et al. 2017a). Acute and sensitive vision supports precision flight by stabilizing scenes during hovering and processing rapid optic flow during forward motion (Sponberg et al. 2015; Stöckl et al. 2017a; Menz et al. 2022).

Hawkmoths have superposition compound eyes. These eyes differ markedly from single lens camera-type eyes, such as those found in vertebrates or spiders. They are made up of arrays of visual units, called ommatidia, that are typically packed in a hexagonal lattice and can be categorized into two kinds, superposition or apposition (Stavenga 2006). Most day-active insects, such as bees and butterflies, possess **apposition** compound eyes, in which each facet focuses light onto a fused bundle of photoreceptor microvilli, the rhabdom (Land and Nilsson 2012).

Light along a facet’s anatomical axis reaches the rhabdom, but off-axis light is absorbed by surrounding screening pigments. One can think of a rhabdom as a pixel (although each rhabdom can contain microvilli from 5–9 photoreceptors) with each facet directing light onto a single spot in an image (Pichaud and Casares 2022). Insects active in dim light, such as moths, beetles, and owlflies, instead possess **superposition** compound eyes, an adaptation that greatly enhances light-gathering ability and sensitivity affording better night vision (Warrant and McIntyre 1990; Warrant 2001; Meyer-Rochow and Gál 2004; Belušič et al. 2013). Unlike apposition eyes, which screen out off-axis light, superposition eyes have a clear zone between lenses and retina, and optics that redirect incoming light from many facets onto a single rhabdom. Optical theory suggests that the increase in sensitivity comes with trade-offs, one of them being that superposition eyes need to be spherical and have equally sized facets across the eye to successfully implement superposition (Warrant et al. 1999; Stavenga 2006)

Moths and butterflies have transitioned between diurnal and nocturnal lifestyles many times in their evolution (Kawahara et al. 2018) and correspondingly they have both apposition and superposition eyes and some intermediate forms (Fischer et al. 2014). The distribution and evolution of superposition and apposition eyes across Lepidoptera is unclear, but studies have proposed links to preferred activity times or diel niche (Horridge et al. 1977; Land 1984; Greiner 2006; Yack et al. 2007; Fischer et al. 2012). However, because diel niche often represents a spectrum rather than a distinct set of categories and can also change in a species (Kirse et al. 2025), disentangling eye and diel-evolution can be difficult.

To understand the relationship between superposition architecture and diel niche, we examined the eyes of six sphingid moth species, three with fully or partial diurnal activity and three with mostly nocturnal activity. We used microcomputed tomography (μCT) to reconstruct the internal and external eye structure, then examined compound eye parameters as they varied across the eye, either from center to periphery or dorsal to ventral, using specialized software (Currea et al. 2023). We estimated local sphericity using radial distance, eye size, facet size, cone skewness and interommatidial angles (Δφ). Crystalline cones are not always perfectly perpendicular to their facet surface; some are skewed, causing the ommatidium to view along a slightly oblique axis. We used ODA to first estimate the anatomical axes of each cone. This allowed us to estimate (1) cone skewness as the angle between a tangent from the eye surface and the cone axis and (2) the Δφ as the mean visual angle between each cone and its six neighbors (Currea et al. 2023). We tested the idea that superposition eyes need to be spherical and examined if diurnal and nocturnal moths both held to this constraint (Stavenga 1979, 2006; Warrant et al. 1999). We also tested if superposition eyes would lack regional specialization or acute zones and examined how acuity, facet size and skewness changed across the eye.

## Methods

### Sampling design and choice of study species

We chose six hawkmoth species from two subfamilies (Macroglossinae and Sphinginae) representing different degrees of diurnality and nocturnality as adults: one species is exclusively diurnal (*Macroglossum stellatarum*), two species are both diurnal and nocturnal (*Macroglossum fritzei, Hyles lineata*), and three species are primarily nocturnal (*Deilephila elpenor, Sphinx ligustri, Manduca sexta*). Diurnality and nocturnality designations were determined based on published literature and consultation with experts. The species selected, along with information on the specimens used, are listed in Supplemental Table 1.

### Microcomputed tomography

To visualize internal eye structure, we conducted μCT scanning at both the Natural History Museum (NHM), London, UK and the University of Florida (UF), Nanoscale Research Facility, Gainesville, FL, USA. Below we outline the protocol followed at each institution.

For specimens imaged at UF, we collected the heads of each species and processed them with a standardized preparation protocol for μCT scanning (Bray et al. 1993; Metscher 2009). Each specimen was initially preserved in a 1.5 ml Eppendorf tube containing 70% ethanol for at least 24 hours. Following preservation, we replaced the ethanol with a 0.3% phosphotungstic acid (PTA) solution prepared in 70% ethanol and allowed specimens to stain for at least three weeks. In most cases, we scanned both eyes, but as some specimens showed collapse on either side, we sometimes only scanned a single eye.

Eyes larger than 3 mm in diameter were removed directly from PTA and scanned in a wet state, (ZEISS Versa 620 XRM). For eyes smaller than 3 mm, dehydration during scanning caused significant shrinkage and displacement, leading to motion blurring. To prevent this, we scanned smaller eyes in a dry state.

To prepare dry samples, we followed a graded ethanol dehydration process. Specimens were sequentially immersed in 80%, 90%, and 100% ethanol, each for 1 hour. Following dehydration, samples were transferred to a 1.5 ml Eppendorf tube containing hexamethyldisilazane (HMDS) for 24 hours, to facilitate infiltration and stabilize internal eye structures. After infiltration, samples were removed from the HMDS solution and placed in a dry, capped Petri dish to allow for slow evaporation. All μCT scans at the University of Florida’s Nanoscale Research Facility were performed using Phoenix vjtomejx M (GE Measurement & Control Solutions) or a ZEISS Xradia 620 Versa 3D X-ray microscope. The ZEISS system is equipped with an automatic reconstruction feature that processes scan data immediately after a complete scan.

For specimens scanned at the NHM, mouthparts and antennae were removed from each head using a dissection microscope to increase the staining efficiency of internal eye structures. The specimens were then fixed in 70% ethanol for at least 24 hours. Following fixation, the ethanol was replaced with 1% PTA solution prepared in 70% ethanol and then allowed to stain for seven days. To examine the effectiveness of staining, specimens were individually scanned. When staining was insufficient, we placed the specimen back into 1% PTA solution and scanned every two days until staining was complete, or for a maximum of 21 days. These μCT scans were performed using a ZEISS Xradia 520 Versa 3D X-ray microscope.

### Scan data processing

After μCT reconstruction, we aligned each scan to a consistent dorso-ventral and anterior-posterior orientation in 3D Slicer (Slicer 5.6.2; Fedorov et al. 2012). We manually extracted ommatidial traces using segmentation tools in 3D Slicer for one eye on each specimen. Segmentations were exported as NRRD files and converted to TIFF stacks via the pynrrd package (See supplementary file 3 for custom code). Raw volumes are available on MorphoSource, and manually traced datasets are on Figshare (https://figshare.com/s/44be67162143d41fddb6).

### Automated measuring of ommatidia with the ommatidia detecting algorithm (ODA)

We automated the analysis of pre-filtered stacks using an established ODA pipeline (Currea et al., 2023). The ODA applies a fast Fourier-transform-based filter to detect the hexagonal lattice of ommatidia in each 2D slice, automatically identifying individual facets and computing their centroid coordinates. For every CT scan, the pipeline outputs two files (ommatidial_data.csv and interommatidial_data.csv, see Supplemental data) with ommatidial counts, center locations, skewness, diameter and radial distances of each crystalline cone. We measured the Δφ of each pair of adjacent ommatidia (Supplemental File: interommatidial_data.csv) and the mean Δφ of each ommatidium and its six neighboring ommatidia (Supplemental File 2: ommatidial_data.csv).

### Defining and measuring eye parameters

#### Radial distance

This is defined as the distance from the center of the best-fitting sphere to the lens centroids fit using a least squares algorithm based on (Jekel et al. 2016) and further described in (Currea et al. 2022).

#### Sphericity

We calculated two putative measures of sphericity. A) 1 - (std. dev(r) /mean(r)), where *r* is the set of all radial distances. This is a complement of the coefficient of variation of eye radial distance across all ommatidia, such that as the standard deviation of radii (normalized by mean radius) approaches 0, this value approaches 1 as it would for a perfect sphere. B) Inner radius/outer radius: we fit two spheres and compare the radius of the minimum bounding and maximum inscribing spheres (Fig. 1).

**Figure 1.**
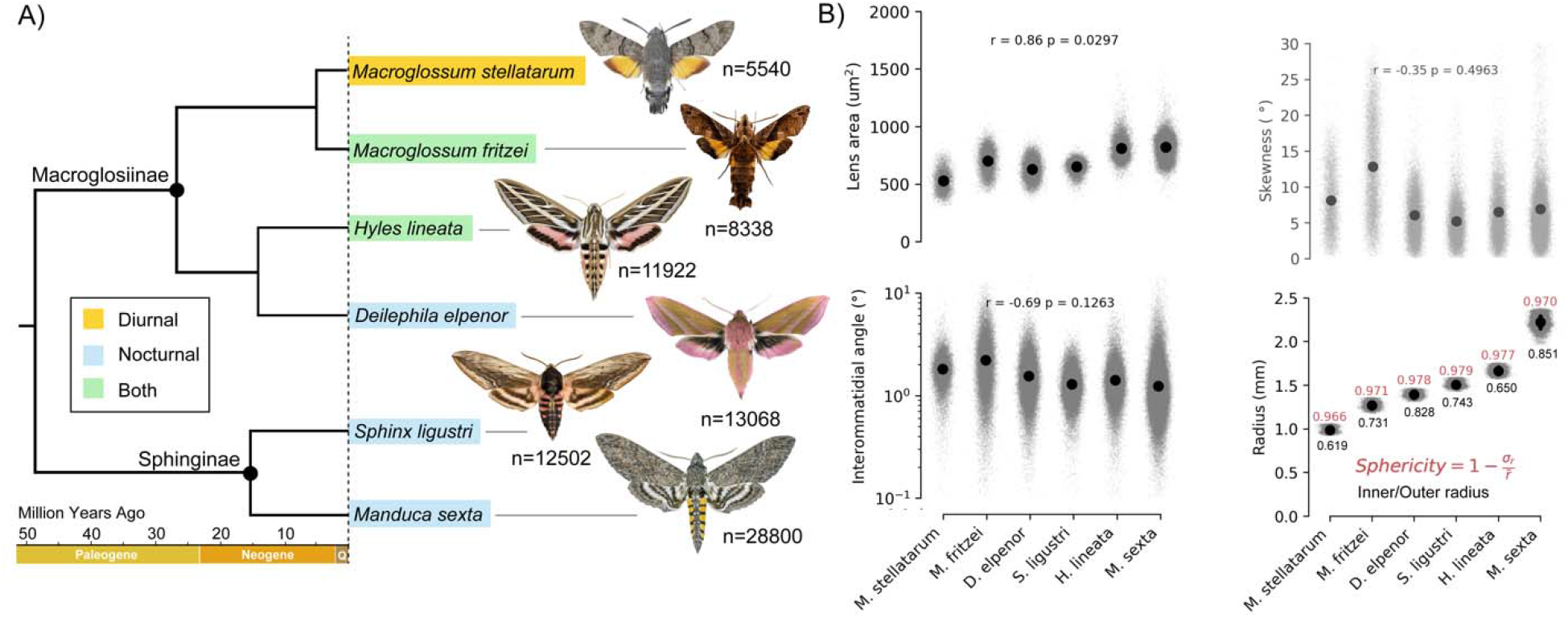
The six species tested for overall ommatidial parameters and eye sizes. A) Phylogenetic tree reconstructed from Kawahara and Barber (2015) and Kawahara et al. (2009). B) Summary statistics for the six species. Small gray dots indicate measurements for individual facets and larger black spots indicate mean values for lens area, IOA, skewness angle, and facet radius along with 99.9% confidence intervals, although these are often so small, they are hidden by the mean markers. Moth species are sorted in ascending order of mean facet radius as an estimate of general eye size. Sphericity—a measure of how close the distribution of radiuses approximates a sphere with 1 indicating a perfect sphere—is printed in red above each distribution of facet radii per species. In black below is a similar metric, inner/outer radius.

Δφ: We calculated Δφ as the mean angular separation between the visual axis of a crystalline cone and those of its six nearest neighbors.

#### Lens diameter

The lens diameter of a given ommatidium was approximated by its mean center-to-center distance between adjacent ommatidia. The lens area of a given ommatidium was approximated by assuming a circular aperture with a diameter equal to the lens diameter. We used the mean radial distance as a measure of eye size. For each species, we altered the algorithm parameters, such as density range and window size, to let the program recognize every ommatidium (see code Supplementary file 3).

#### Cone skewness

Skewness for each ommatidia is the tilt of cone axes, relative to the local surface normal. It was calculated as the angular deviation of the cone axis from the local facet normal, which is equivalent to 90° minus the angle to the tangent plane (Stavenga 2006).

### Manual measuring of ommatidia

To validate some of the automated measurements, we manually measured the facet or lens diameter and (Δ□) along three axes passing through the center of the eye: equatorial, 120° above, and 120° below the equator. Along each axis, we sampled five regions, each spaced by ∼15° of viewing angle. We made the measurements in 3D Slicer on the 2D cross-section panel oriented to the tested axis. In each sampled region, we measured the lens diameter by taking the average end to end width of three successive facets. For the Δφ, we traced back 5 ommatidia to the point of intersection and divided the angle by 5. We also estimated the eye size of each species by measuring the diameter of the eye horizontally.

### Capturing representative 2D images

After reorienting each scan in 3D Slicer, we generated two orthogonal cross-sections passing through the estimated geometric center of each eye. One section was in the transverse plane (horizontal to the eye) and the other in the sagittal plane (vertical to the eye).

### Statistical analyses and measuring eye parameters

We calculated the mean and 99.9% confidence intervals of the mean for lens area, Δφ, skewness angle, and radial distance of the facets and then measured the Pearson correlation of lens area, Δφ, and cone skewness relative to mean eye radius. Parameter definitions and computational details follow Currea et al. (2023); briefly, we measured lens diameter, lens area, Δφ, cone skewness, and radius in both transverse and sagittal planes. Using ordinary least squares regression, we modeled lens area, which limits optical sensitivity, as an affine function of skewness angle.

## Results

We successfully constructed μCT scans of the six hawkmoth species (Fig. 1). We then oriented all reconstructions in the same direction, cleaned and cropped each to a single eye, and manually cleaned until the crystalline cones became visible (Fig. 2, S1, S6). We then estimated all eye parameters with the ODA and manually aligned the final maps with the CT scans to identify the dorso-ventral and antero-posterior axes (Fig. 3)

**Figure 2.**
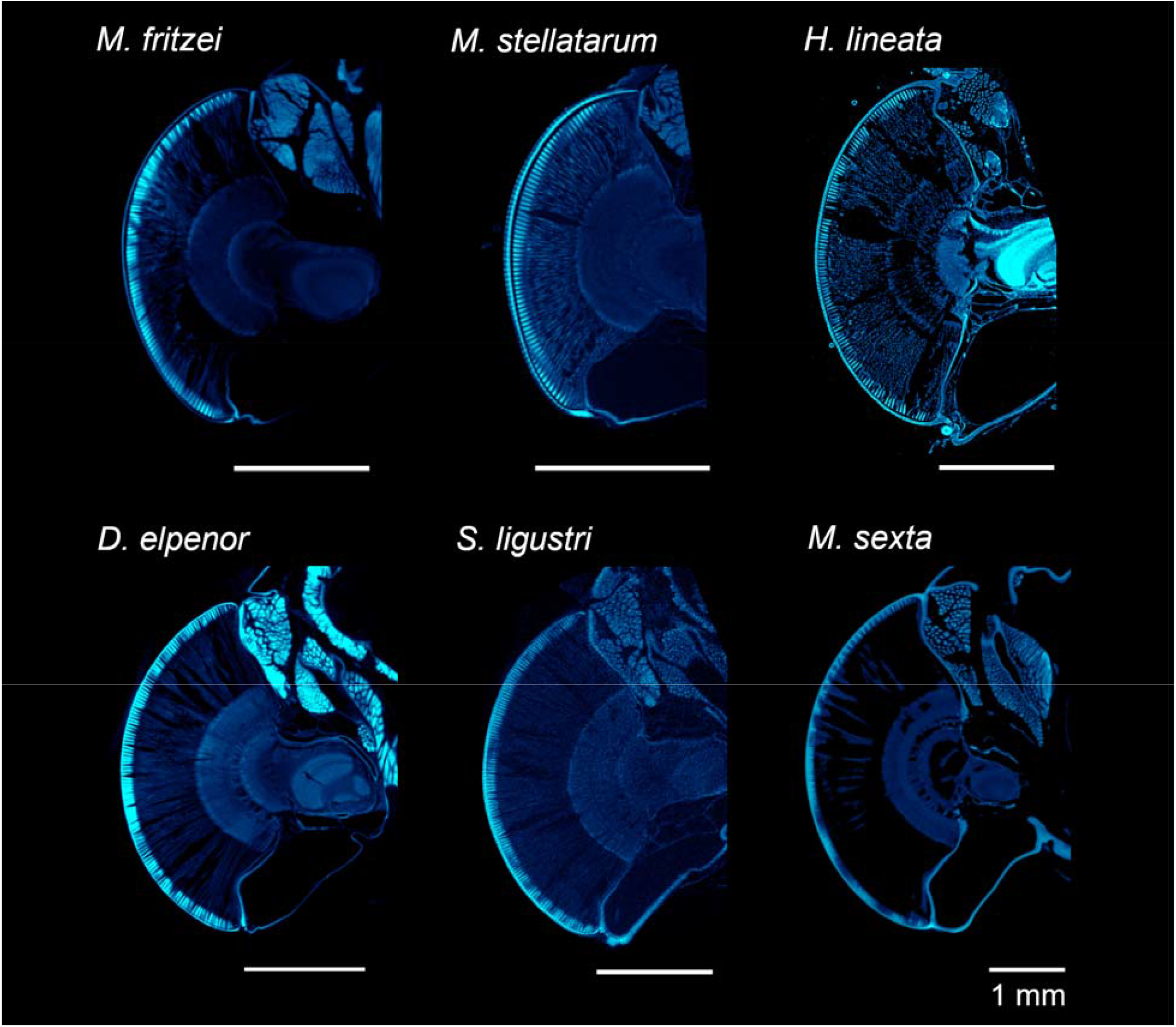
Transverse μCT scan cross-sections of the compound eye of the six hawkmoth species studied. Each cross-section passes through the center of the eye. In all images, the top corresponds to the anterior side of the head, and the bottom to the posterior.

**Figure 3.**
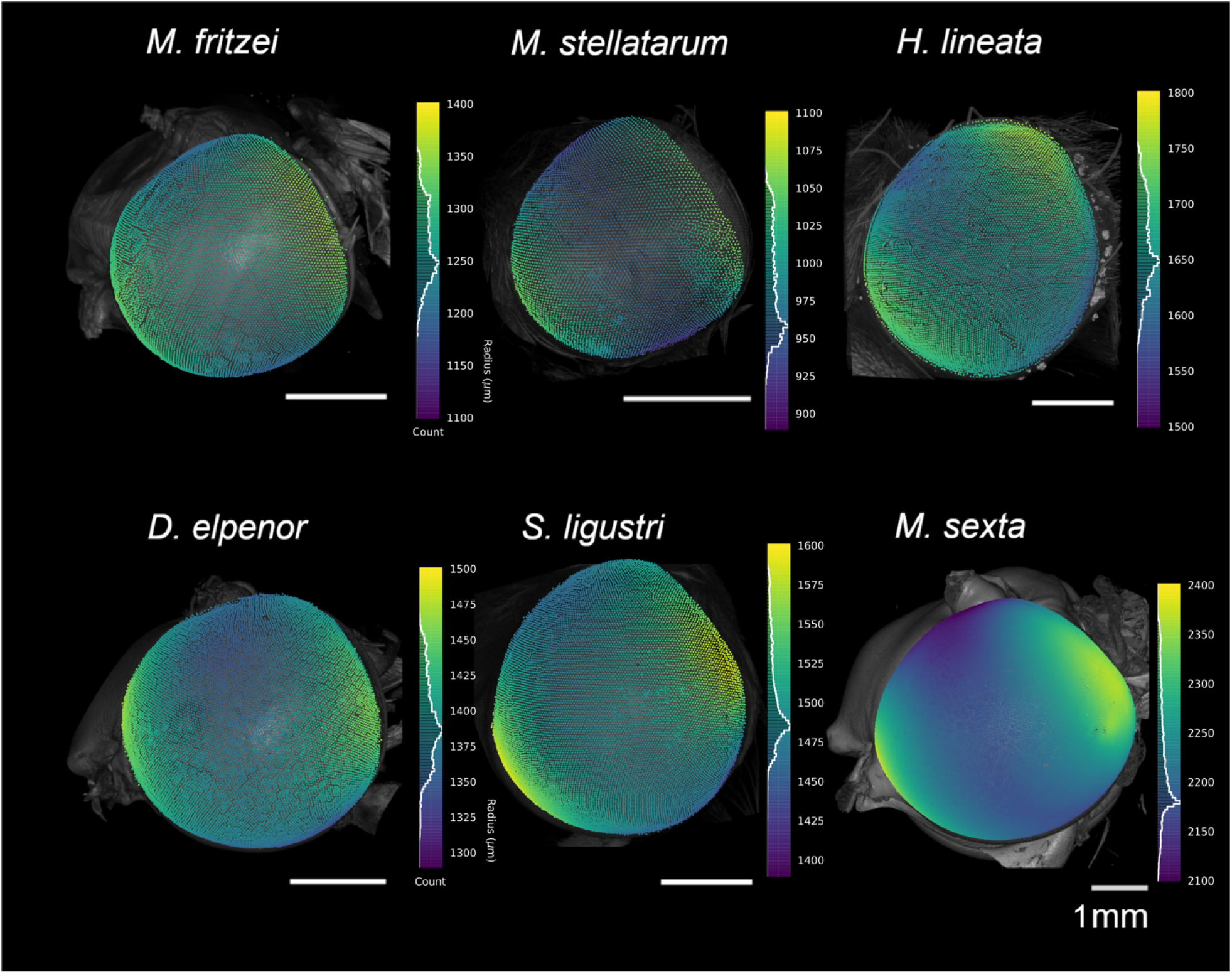
Variation in estimated radius (sphericity) mapped onto the 3D surface of the eye to showcase variation from center to dorsoventral (decrease) and anteroposterior edges (increase). The histograms on the right indicate counts for the estimated radius values at various ommatidia. Data were rotated to a global coordinate orientation.

### Ommatidial counts, facet size, Δφ and correlations with eye size

We first quantified the number of ommatidia and variation in lens area across species. Counts ranged from 5,500 to 28,800 ommatidia (Table 1, Fig 1A) and increased with eye diameter (r = 0.95, p = 0.036; Fig. 1B). The diurnal species *Macroglossum stellatarum* had the smallest eyes and lowest ommatidial counts (∼5,540), while the crepuscular *Manduca sexta* had the largest eyes and highest counts (∼28,800). Facet size (lens area) was positively correlated with eye size (r= 0.86, p=0.029, Fig. 1B), with the smallest in *Macroglossum stellatarum* and largest in *Manduca sexta*. Δφ was negatively correlated with eye size (r = -0.69, p =0.13, Fig 1B), with *Manduca sexta* exhibiting the lowest Δφ (highest spatial acuity).

**Table 1.** List of species analyzed in this study. Parameters used for cleaning and the range of ODA output values for single eyes are listed. (D: Diurnal, N: Nocturnal)

| Species | Diel Niche (D/N) | Specimen No. | Sex | Eye analyzed (R/L/Both) | CT Voxel (μm) | Facet counts | Facet diameter (μm) | Δφ | Eye diameter (mm) |
| --- | --- | --- | --- | --- | --- | --- | --- | --- | --- |
| *Manduca sexta* | N | 13460 | - | Left | 4.50074 | 28800 | 32.29±1.77 | 1.24±0.81 | 4.426 |
| *Deilephila elpenor* | N | Bb_02 | Male | Right | 3.325 | 13068 | 28.21±1.86 | 1.55±0.75 | 2.754 |
| *Hyles lineata* | D/N | HL002 | Male | Right | 3.4784 | 11922 | 32.05±1.85 | 1.41± | 2.967 |
| *Macroglossum fritzei* | D/N | LEP29152 | Male | Right | 5.6845 | 8338 | 29.81±2.21 | 2.22 ± 1.40 | 2.452 |
| *Macroglossum stellatarum* | D | M002 | Female | Right | 2.4 | 5540 | 25.85±2.16 | 1.82±0.63 | 1.906 |
| *Sphinx ligustri* | N | SLF02 | Female | Right | 3.3 | 12502 | 28.80±1.10 | 1.28±0.43 | 2.836 |
| *Sphinx ligustri* | N | BB_09 | Male | Both | 4.7828 | 10955 | NA | NA | 2.937 |

### Eyes are close to spherical, but have consistent local changes in radial distance

We measured eye size using automated fitting of ommatidial centers to a spherical surface (Table 1) and manual measurements of eye diameter (Table S2). Radial distance varied systematically across the eye surface (Fig. 3), revealing consistent local departures from a perfect sphere. We found that in all six species, the eye was flatter along the antero-posterior axes and more curved along the dorso-ventral axis (Fig 3, S7). We estimated global sphericity using the coefficient of variation of radial distance and found that all species eyes had values close to 1, but when fitting a minimum bounding and maximum inscribing sphere (inner/outer radius) the diurnal species showed more deviations from a perfect sphere than nocturnal species (Fig 1B, See methods for more details).

### Δφ increases from center to periphery in most nocturnal species

Δφ varied systematically across the eye axes (Figs. 4, S2, S3). In all species, Δφ was lowest in the central region of the eye and increased towards the periphery (Fig. 4, Table S2). Local irregularities in Δφ values were visible across the eye surface in some species (Fig. 4), but these did not persist in the binned means (Fig. 4, S3). Among the six species, *Macroglossum fritzei* showed the most abrupt shifts in Δφ, whereas the remaining species showed more gradual changes.

**Figure 4.**
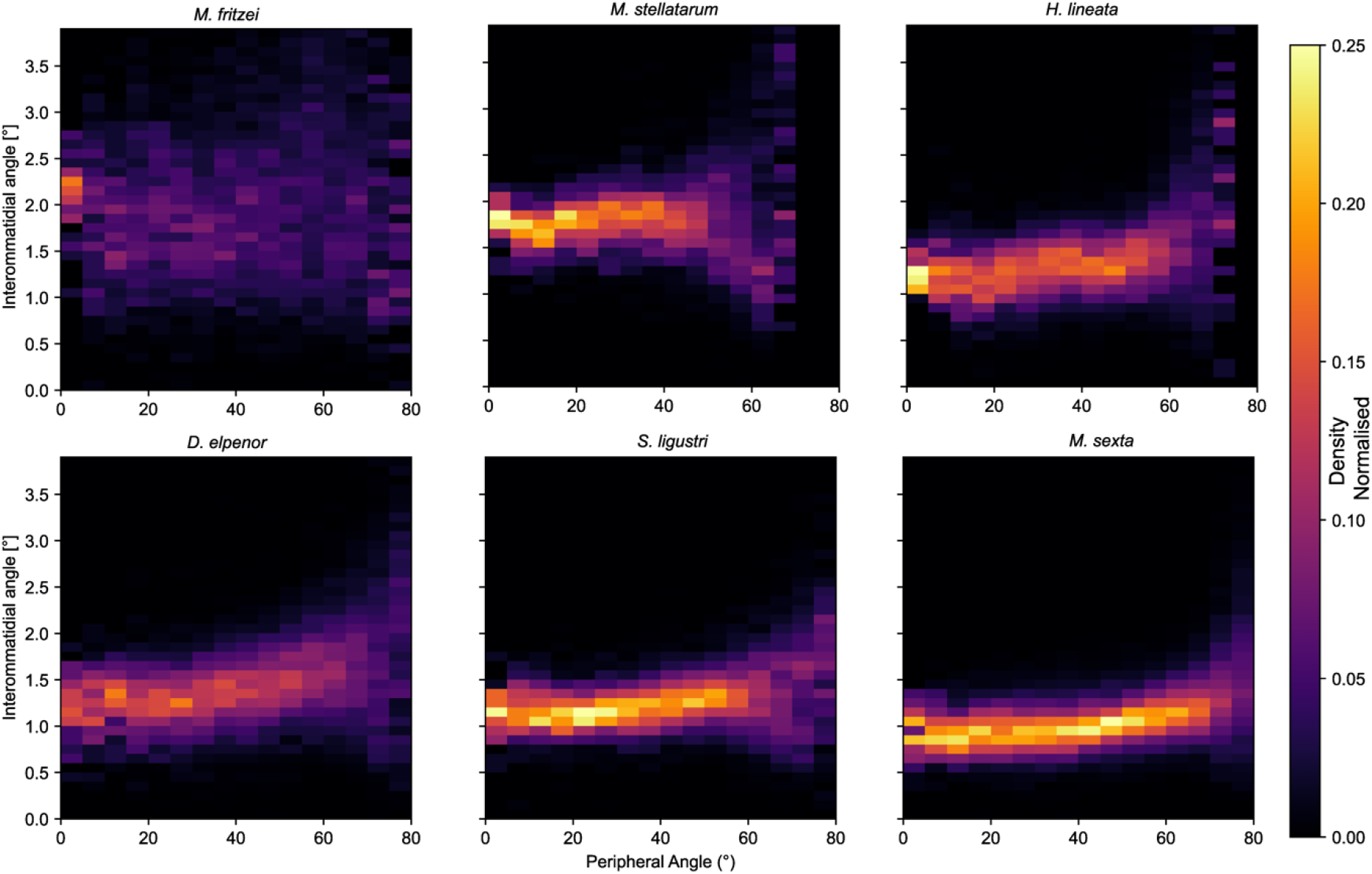
Variation in Δφ from center to periphery. Species 1-3 (top row) have some diurnal activity and the three species on the bottom are nocturnal, axes and data has been truncated to showcase most variation and to make plots comparable.

### Facet size increases and decreases gradually from center to periphery in different species

Lens diameter and area varied systematically across eye axes, but the pattern was not consistent across species (Figs. 1B, Fig 5, S4, S5). We collapsed variation onto a single peripheral ring (eccentricity from 0–90°, Fig. 5) and found that some species show clearer trends: *Manduca sexta* exhibited a ∼15–18% decrease in facet diameter from center to periphery, whereas both *Macroglossum* species showed steeper declines (∼20–23%). In all tested nocturnal species, we observed only gradual decrease from center to periphery along both the dorsoventral and anteroposterior axes (Fig. S5) confirmed with manual measurements (Table S2). Notably, *Macroglossum stellatarum* and *Macroglossum fritzei* had a distinct large facet-size zone in the dorsoposterior region, clearly visible in the eye maps and the binned means (Figs. S4).

**Figure 5.**
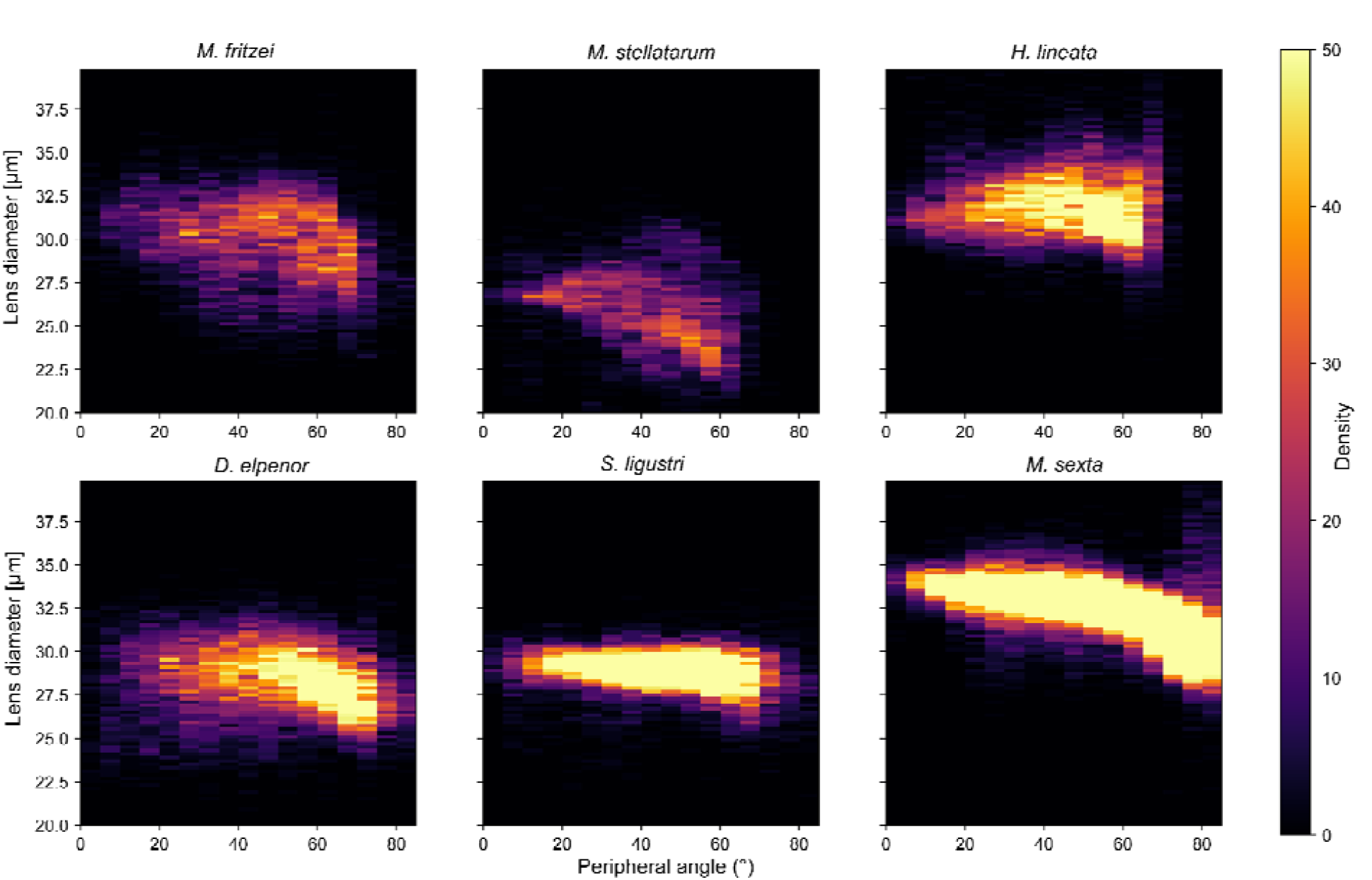
Comparison of diel-activity change in relation to eye diameter and peripheral angle. Peripheral angle was a measure of center or periphery, defined as where theta and phi refer to the polar representations of the ommatidia. Species 1-3 (top row) exhibit diurnal activity, and the three species on the bottom row are nocturnal. Species depicted (from upper left to bottom right)

### Nocturnal species show minimal cone skew

All three nocturnal species exhibited minimal skew across both axes (Fig. 6). In contrast, *Macroglossum fritzei* and *Macroglossum stellatarum* displayed a sharp increase in skew toward both the posterior and anterior regions of the eye, exceeding 20 degrees in the posterior of *Mac. fritzei* (Fig 1B). *Hyles lineata* showed a similar but smaller increase towards the posterior eye. Using ordinary least squares regression, we modeled lens area, as an affine function of skewness angle. We found significant positive correlations between skew angle and lens area in the 4 Macroglossiinae (r > 0.18, p < 1×10^-57^) but not in the 2 Sphinginae (r < 0.002, p > 0.73; Table S3). Lens area increased by 4.1 to 4.8 μm^2^ for every degree increase in skewness angle, as opposed to 0.02 to 0.03 μm^2^/deg in *S. ligustri* and *M. sexta*.

**Figure 6.**
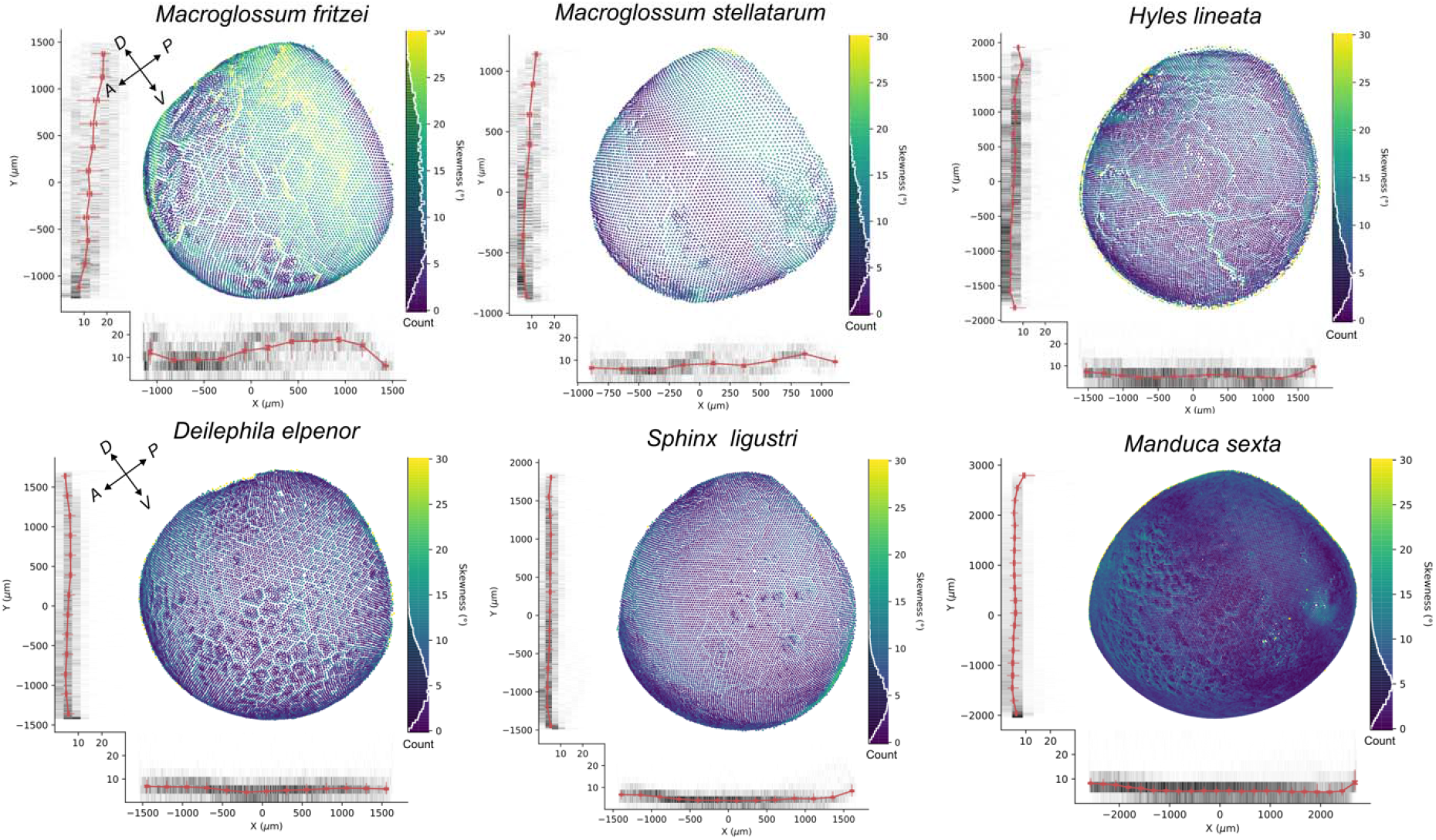
Variation in cone skewness in X (posterior to anterior) and Y (ventral to dorsal) for all the hawkmoth species studied. Red lines depict individual counts along the X and Y axes. Data were rotated to a global coordinate orientation.

## Discussion

### Optical theory of superposition eyes

In a dark-adapted superposition eye, each rhabdom receives light from many corneal facets, which greatly increases sensitivity. But this requires that each facet, in turn, correctly directs light towards many rhabdoms, depending on angle of incidence, and this significantly constrains the eye’s geometry. This implies superposition eyes might lack the regional specializations that characterize many apposition eyes, as the surface may need to remain nearly spherical, with facets of relatively uniform spacing and size. Our results generally support this view. Across six sphingids displaying both diurnal and nocturnal activity, we found their eyes remaining close to spherical. Δφ varied gradually rather than forming discrete acute zones, and facet sizes changed smoothly across the eye surface. However, we did find repeatable deviations from perfect spherical symmetry. All tested species showed flattening in the antero–posterior plane and greater curvature in the dorso–ventral plane, indicating that superposition optics tolerate modest regional distortion. Further, diurnal species showed stronger local departures, including pronounced crystalline cone skew in posterior regions of *Macroglossum*. This suggests superposition eyes are not rigidly constrained to perfect spherical geometry but operate within an envelope which can compensate for moderate distortions—potentially through cone skew or subtle changes in facet geometry. So rather than prohibiting regional specialization, superposition architecture may simply limit its magnitude and sharpness.

### Diel-evolution and previous visual ecology studies in hawkmoths

Although most hawkmoths are night-flying, some are strictly diurnal and others show both day and night activity (Beck and Linsenmair 2006; Broadhead et al. 2017). Within hawkmoths, diurnal reversions have occurred independently in several genera, sometimes involving only a small number of species across three of the four subfamilies. These include *Afrosatspes* and *Sataspes* (subfamily Smerinthinae), one species of *Sphinx* (subfamily Sphinginae), and *Cephonodes* and *Hemaris, Aellopos, Euproserpinus* and most *Proserpinus*, as well as *Hayesiana* and some species of *Macroglossum* (subfamily Macroglossinae). However, within these diurnal species, vision has only been studied extensively in the hummingbird hawkmoth, *Macroglossum stellatarum*. In this species, the superposition pupil (the effective superposition aperture visible as the eye glow) remains unusually open even in bright light, so the eye retains a refracting superposition state under conditions in which many moths would reduce optical pooling through pigment migration (Warrant et al. 1999; Stöckl et al. 2016, 2022). Relative to more nocturnal hawkmoths, however, the superposition aperture is smaller, consistent with adaptation to brighter visual environments (Stöckl et al. 2017b). No other diurnal hawkmoths superposition architecture has been studied in similar detail, making it difficult to determine whether these features are exceptional or a pattern present across all diurnal hawkmoths. To explore how eye architecture relates to diel niche, we therefore selected a mix of diurnal and nocturnal species, prioritizing species with known visual ecology, or those with accessible colonies, and explored using 3D eye scans.

### Hawkmoth eyes are near-spherical in shape, with axis-specific departures

Quantifying sphericity in biological eyes is inherently metric-dependent, because “sphericity” is not uniquely captured by any single index. We therefore report two complementary metrics (globally and locally) as descriptive summaries of eye-shape variation. Results from both metrics suggest minimal deviation from a spherical eye shape. *Macroglossum stellatarum* exhibits the lowest sphericity among the species examined, but still closely approximates a sphere—especially compared to the much less spherical eyes in many other insects.

Within each eye, we found consistent variation in estimated curvature radius in both eye axes dorso-ventrally and antero-posteriorly across all species, providing a measure of how flat or curved the eye is at different regions. This suggests that all species show regions of local flattening and regions with more curvature. Interestingly, the orientation of this curvature variation was not consistent across species, not even between the different *Macroglossum* species. This suggests that species may be optimizing the balance between field of view (more curved) and higher acuity (flatter), in ways that match their flight and motion requirements. This warrants further exploration in future studies, as it could reflect a developmental constraint as much as an adaptive benefit.

### Nocturnal species have larger eyes featuring larger and more numerous facets than the strictly diurnal species

Our results show that strictly diurnal species have the smallest eyes, with the fewest ommatidia and smallest facet size (Table 1; Fig. 1B). Across the six species, eye size was positively correlated with both facet count and facet size (Fig. 1B), consistent with the expectation that larger eyes accommodate more sampling units and larger optical apertures. We did not find evidence that strictly nocturnal species necessarily have the largest eyes in this dataset, because *Hyles lineata*, which is active both during the day and at night, had the second-largest eye size and ommatidial count. Our results are consistent with results published by (Yagi and Koyama 1963), which reported similar scaling between eye size, facet number, and facet dimensions in Lepidoptera.

In insects with apposition eyes, larger facets in nocturnal taxa are commonly interpreted as an adaptation for increased photon capture and thus higher optical sensitivity in dim light (Somanathan et al. 2009). Although superposition eyes differ in that sensitivity is strongly shaped by optical pooling and by pupil dynamics, facet size and eye size should still contribute to potential light capture when the superposition pupil is open. We did not quantify pupil size under fully dark-adapted conditions in our specimens, so we cannot estimate maximal optical sensitivity directly from our scans. Nevertheless, the fact that nocturnal species retain larger facets and larger eyes even in the light-adapted state suggests that these differences are not solely a transient consequence of pupil state. Our results are consistent with the interpretation in previous studies that nocturnal hawkmoths achieve higher potential sensitivity through a combination of larger optical apertures and a larger pupil. (Yagi and Koyama,1963; Stöckl et al. 2017b)

### Intraocular variations in facet size and Δφ

Both facet size and Δφ vary across the eye surface within each species, and these intraocular gradients tend to be steeper in the diurnal species. We previously hypothesized that diurnal taxa might show more abrupt regional changes in ommatidial size and spacing, and the strictly diurnal *Macroglossum stellatarum* provides some support for this idea: despite having the lowest facet counts, it shows the sharpest within-eye transitions in both facet size and Δφ, consistent with previous reports (Warrant et al. 1999; Stöckl et al. 2022). However, we did not find evidence that nocturnal species lack regional variation across the eye; instead, they also show regional change, but typically with more gradual transitions. One possible explanation is that diurnal hawkmoths may operate their superposition eyes under light-adapted conditions more frequently, when effective optical pooling is reduced and regional differences in ommatidial size and spacing may have stronger functional consequences. This idea remains speculative and will require future tests linking pupil state, light adaptation, and spatial variation in optical performance.

Patterns of facet-size variation differed among species. *Manduca sexta* showed an antero-ventral region with larger lenses that decreased in size both ventrally and toward the periphery. *Sphinx ligustri* and *Deilephila elpenor* showed comparatively the least changes in facet size across the eye. *Hyles lineata* showed an entirely different pattern with an almost anterior dorsal bump in facet size. Taken together, these results indicate that both diurnal and nocturnal taxa can exhibit pronounced regional tuning of facet size, but the form of that tuning is species-specific rather than neatly partitioned by diel niche.

In contrast to facet size, Δφ was relatively uniform across the eye surface in all species, with most species showing a modest increase toward the periphery (Fig 8-9). Despite the center of the eye generally providing the highest spatial resolution (lowest Δφ), none of the six species exhibited a distinct acute zone in the sense of a sharply bounded region with strongly reduced Δφ. Instead, acuity changes were gradual, producing a broad central region of slightly enhanced resolution rather than a pronounced fovea-like specialization. This pattern is expected because coherent imaging in superposition eyes depends on maintaining near-spherical curvature, and therefore major local departures such as localized flattening that would strongly compress Δφ are limited. Taken together, our facet size and Δφ maps and curvature measurements support the hypothesis that superposition architecture favors relatively uniform sampling with only modest, gradual regional tuning, rather than highly localized acute zones.

### Cone skewness as a regional specialization in tribe Macroglossinae and genus Macroglossum

Compared with the other species, *Macroglossum stellatarum* and *Macroglossum fritzei* showed a pronounced posterior region of highly skewed crystalline cones, consistent with previous reports from the former species (Warrant et al. 1999). This pattern is only evident in antero–posterior views but not in dorso–ventral sections, suggesting that the skewness is directionally structured rather than a global tilt expressed across all axes. A more plausible geometric interpretation is that the pronounced posterior skewness redistributes ommatidial viewing directions and likely expands posterior field of view. This tradeoff would be expected to increase local Δφ along the antero-posterior axis and therefore reduce both acuity and sensitivity in the affected region. *Macroglossum* species are exceptionally fast and maneuverable, and a regional visual specialization of this kind could support accurate flower tracking with frontal vision while enabling rapid responses to peripheral motion during evasive flight. No other species in our sample shows comparably strong posterior skewness, suggesting this may be a *Macroglossum* associated adaptation rather than a general feature of Sphingidae superposition eyes

Cone skewness is also expected to involve optical tradeoffs. Changing cone axis orientation can redistribute viewing directions and potentially expand regional field of view coverage, but it can reduce sensitivity by decreasing the effective aperture and optical gain of the affected ommatidia. In this context, pronounced cone skewness may be most feasible in taxa that routinely operate under bright or intermediate light levels, where photon limitation is weak and some loss of sensitivity can be tolerated in exchange for altered visual sampling. This provides a possible explanation for why strong skewness is expressed in the diurnal *Macroglossum* species but is absent from the strictly nocturnal species in this study.

Interestingly, we saw a phylogenetic constraint in patterns of cone skewness and lens area, with significant positive correlations between skew angle and lens area in the 4 Macroglossiinae (a weak correlation ∼0.2) but completely missing in the 2 Sphinginae (Table S3). No other metric showed any clear phylogenetic patterns, if anything eye parameters seem to switch rapidly even within a genus, so exploring evolutionary variation of skewness across hawkmoths is good avenue to explore.

Together, these patterns suggest that, within the global geometric constraints of superposition optics, regional tuning may be achieved through coordinated variation in cone orientation and lens size, rather than through a sharply bounded acute zone.

### Patterns of facet and angle distribution: Lepidoptera and other insects

The two central ideas we wanted to test were constraints on variation in superposition eyes and how eye shape and curvature might change. To our knowledge, eye curvature has been examined only by del Portillo (1936), who used histological sections from a few species and found that largely the superposition eyes had equal radii of curvature in both horizontal and dorsal planes and gradual changes in Δφ across the eyes. He examined a large Saturniidae, *Hyalophora cecropia* (then known as *Samia cecropia* or the Cecropia silkmoth), a medium sized Arctiinae, *Euchelia jacobea* (then known as *Tyria jacobaeae* or the cinnabar moth) and a medium sized Notodontidae moth, *Phalera bucephala* (the buff-tip moth) (del Portillo 1936). He found that the *Hyalophora cecropia* eye is equally curved in both median planes, and had its curvature radii homogeneous at all points, except for a small dorsal apex. The same holds for the angular values (presumably Δφ), which are virtually equivalent across most regions; expect at the upper end the frontal section has slightly more wide-angled ommatidia that are inclined toward the corneal surface. In *Phalera bucephala*, the cross section shows less perfect regularity than in *Hyalophora*, but the overall patterns are the same in both species: angular values (Δφ) coincide in the eye’s center and increase slightly forward and downward. In *Euchelia jacobaea*, the eyes are small and nearly spherical, with highly uniform angles and radii; but only the upward-directed ommatidia have slightly wider angles (del Portillo 1936).

Yagi and Koyama (1963) conducted a large-scale survey of facet variation across more than 30 butterfly and moths species and described results similar to ours – a gradual drop in ommatidial size from center to periphery, as well as a positive correlation between facet size and eye size (Yagi and Koyama 1963). They also present facet count estimates for several Sphingidae, including an estimate of 27,000 for *Agrius convolvuli* (Yagi and Koyama 1963). Research on the highly sexually dimorphic moth species *Orgyia antiqua* (Erebidae: Lymantriinae) revealed that females have a lower resolution (Δφ = 3° in females, versus 1.9° in males) and less curved eyes (female = 327 μm and male = male = 461 μm), which is likely due to the fact that females are flightless (Lau and Meyer-Rochow 2007). Warrant et al. (1999) explored eye and facet parameter variation in *Macroglossum stellatarum* in great depth. However, they concluded that optical pooling and rhabdom packing density are more responsible for resolution changes than variation in facet size alone. They did find a posterior region with more skewed cones and suggested this arrangement may maintain resolution needs in the frontal eye, but increase field of view in the posterior (Warrant et al. 1999). Recent work using μCT scans has further examined allometry and individual variation (Stöckl et al. 2022). Other moth eye research compared the angular spread of eyeshine with electrophysiologically measured acceptance angles, demonstrating that moths can vary considerably in whether their visual systems are over or under sampling (Horridge et al. 1977). There are several groups who have studied this in butterflies, such as Pieridae ((Ribi 1979) and Nymphalidae (Rutowski et al. 2009), but since these are largely diurnal and have apposition eyes, more facet variation is expected (Seymoure et al. 2015).

## Conclusion

Using high-resolution μCT and automated whole-eye mapping across six hawkmoth species, we show that hawkmoth superposition eyes are neither perfect spheres nor highly regionalized in the way often seen in apposition eyes. Instead, the eyes remained broadly near-spherical and lacked sharp acute zones, yet still exhibited modest regional tuning in curvature, facet size, and cone orientation, especially in diurnal *Macroglossum*. Although our dataset is limited by low within-species sampling and uses morphological inference alone, it provides a comparative foundation for understanding how superposition eyes evolve across diel niches and for testing how local eye geometry shapes visual function.

## Acknowledgments

We thank Anna Stöckl for providing specimens of *Macroglossum stellatarum* and for comments and discussions. We thank R. Keating Godfrey for providing specimens of the *Hyles lineata* moths. Gary Scheiffele and the staff at the Nanoscale Research Facility at UF helped facilitate our research. Jaime Gray provided resources and assisted with CT reconstruction software, Kelly Dexter, Amanda Markee and David Plotkin provided lab logistical support. We thank

Sönke Johnsen, Victoria Wood and David Plotkin for input on the manuscript. We also thank Alex Ball and the staff of the Imaging and Analysis Centre of the NHM for CT lab and logistical support.

## Author contributions

**Yash Sondhi**: Conceptualization, Investigation, Methodology, Formal Analysis, Writing – Original Draft

**Ruchao Qian**: Conceptualization, Investigation, Methodology, Formal Analysis, Investigation, Writing – Original Draft

**Isabella Koushiar**: Data curation, Writing – Review & Editing

**Ian Kitching**: Conceptualization, Funding Acquisition, Supervision, Resources, Writing – Review & Editing

**John Pablo Currea**: Conceptualization, Software, Formal Analysis, Writing – Review & Editing

**Deborah Glass**: Conceptualization, Investigation, Data Curation, Resources, Writing – Original Draft

**Jacqueline Degen**: Resources, Writing – Review & Editing

**Ed Stanley**: Investigation, Data Curation, Resources

**Simon Sponberg**: Supervision, Funding Acquisition, Writing – Review & Editing

**Akito Kawahara**: Conceptualization, Funding Acquisition, Supervision, Resources, Writing – Review & Editing

**Jamie Theobald**: Funding, Conceptualization, Funding Acquisition, Supervision, Resources, Writing – Review & Editing

## Tables

## Supplemental Files

See link for supplementary data which includes figures, tables and protocols.

## Data Availability

Count data and ODA files are provided on

Figshare: https://figshare.com/s/44be67162143d41fddb6.

The μCT scan for all six hawkmoth species analyzed in this study are publicly available on MorphoSource under the following DOI: 10.17602/M2/M780271; 10.17602/M2/M780277; 10.17602/M2/M780361; 10.17602/M2/M780378; 10.17602/M2/M780384; 10.17602/M2/M780405.

