## Supplemental Information for "Superposition eyes of diurnal and nocturnal hawkmoths are imperfect spheres and lack acute zones"

**Journal:**

**DOI:**

**Corresponding Author:** Yash Sondhi

**This PDF file includes:**

Supplementary Data on Figshare and Morphosource

Supplementary Figures 1 to 7
Supplementary Tables 1-3
Supplementary Methods

Supplementary References

**Other supporting materials for this manuscript include the following:**

The µCT scan for all six hawkmoth species analyzed in this study are publicly available on MorphoSource under the following DOI: 10.17602/M2/M780271; 10.17602/M2/M780277; 10.17602/M2/M780361; 10.17602/M2/M780378; 10.17602/M2/M780384; 10.17602/M2/M780405.

Datasets are: <https://figshare.com/s/44be67162143d41fddb6> : Contains cleaned stacks fed as input into ODA, outputs of ODA after correcting for rotation, previous versions of ODA and some scripts used to make the plots along with ipython notebooks that have the the data and plots already stored.

Supplemental Table 1. Taxon table, status of scan. Table_S1: CT_eye_list

Supplemental File 2: Metadata describing output from ODA for six species.

Supplemental Folder 3: ODA files Output of each species interommatidial.csv and ommatidial.csv

Supplemental Folder 4: Analysis_tools: Analysis code in a python notebook and scripts that describe parameters used for each species (Add to gituhb).

Supplemental file 5: Raw_CT_scans_hawkmoth Tiff stacks of insects zipped

Supplemental file 6: Cross-sections_and_screenshots.zip Zipped file of all the dorsal and ventral cross sections of the moth.

Supplemental Figures

**Figure S1**:

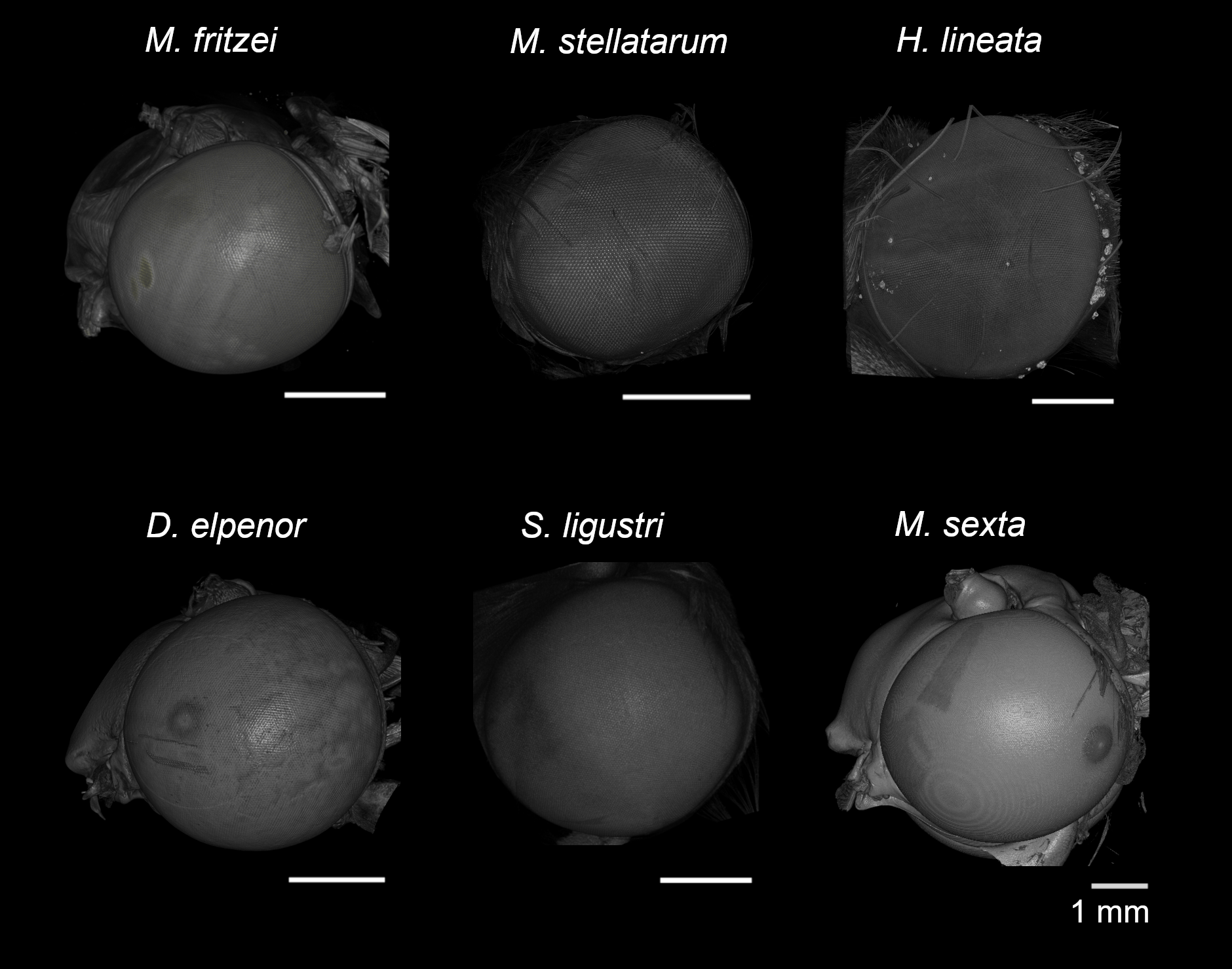

**Figure S1**: 3D lateral image of the compound eye of the six moth species studied. Anterior is to the left, posterior to the right, dorsal at the top, and ventral at the bottom.

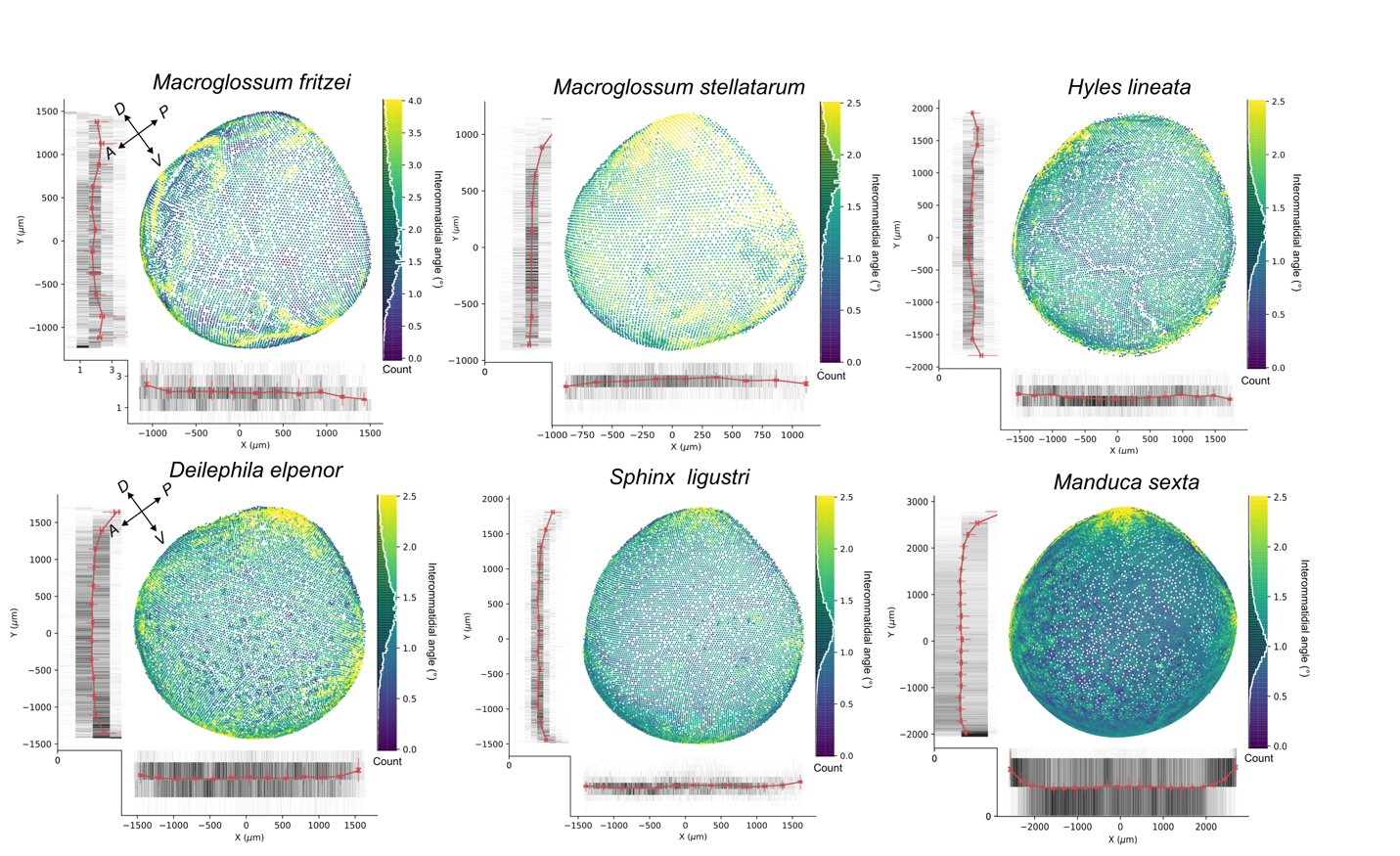

**Fig S2**: Variation in Δφ in X (posterior to anterior) and Y (ventral to dorsal) for all the hawkmoth species studied. IOA distributions were truncated max of 2.5-4 degrees to showcase most of the variation. Red lines depict individual IOA counts along the X and y axes. Data was rotated to a global coordinate orientation.

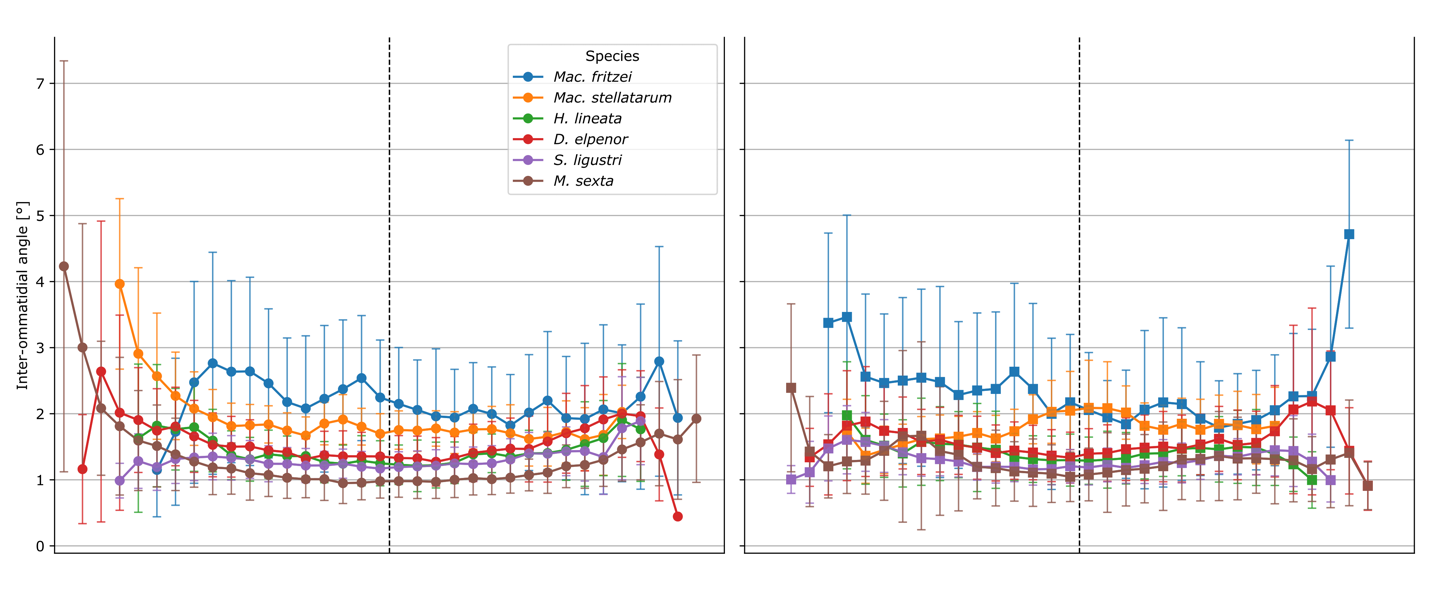

**Fig S3 :** Equatorial (left) and transverse variation (right) in Δφ taken along a single plane for each of the six hawk moth species measured.

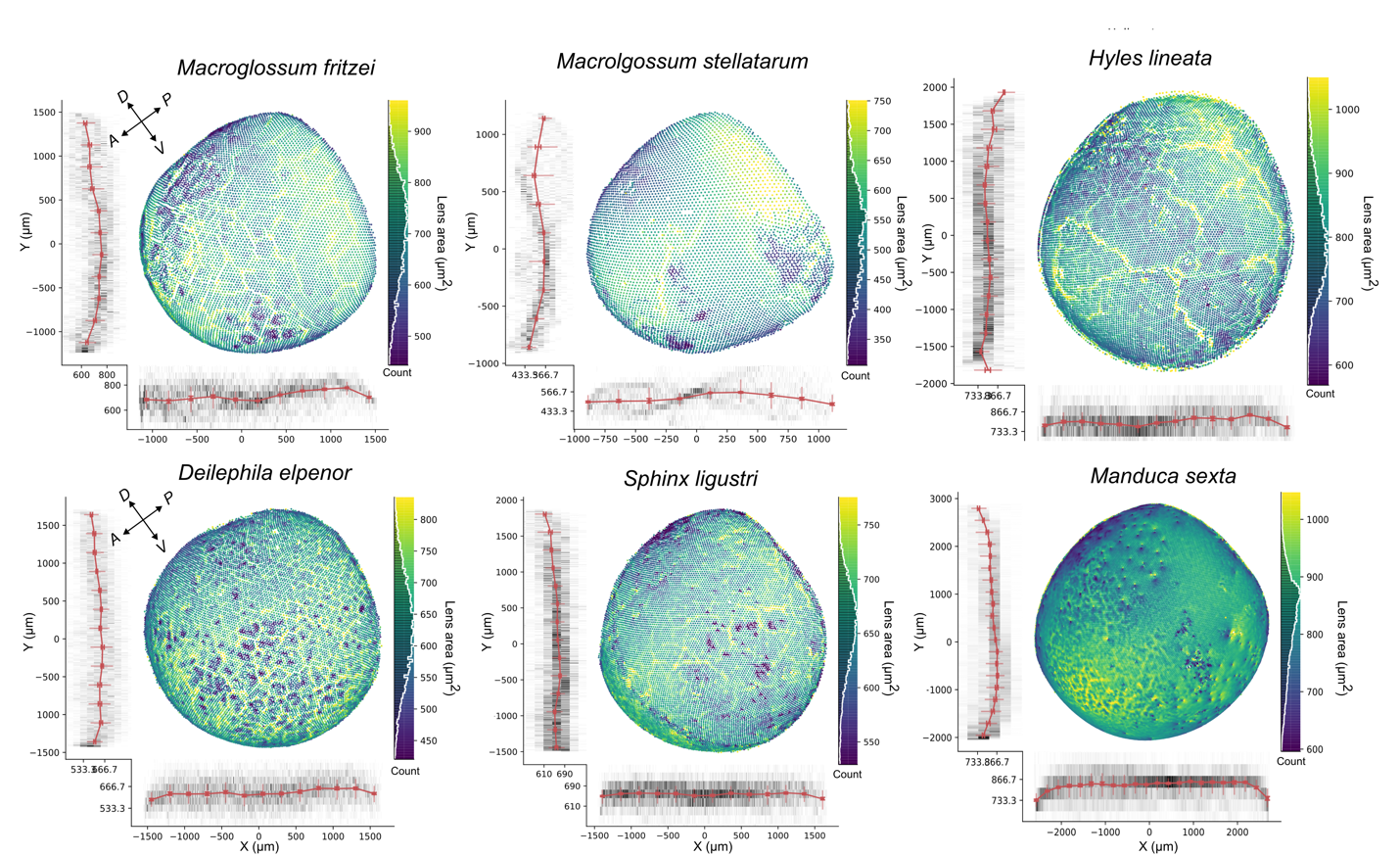

**Fig S4 :** Variation in facet size (lens area), in X (posterior to anterior) and Y (ventral to dorsal) for all the hawkmoth species studied. Red lines depict individual counts along the X and Y axes. Data were rotated to a global coordinate orientation.

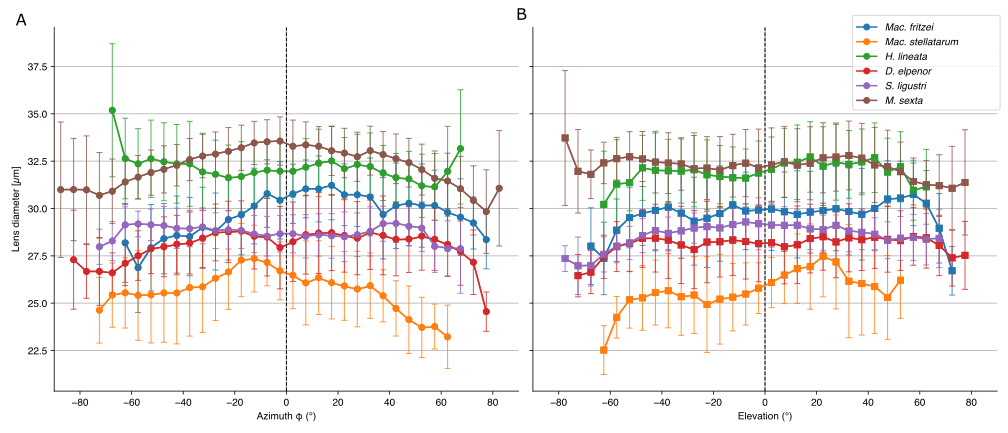

**Fig S5:** Variation in lens diameter in azimuth (left) and elevation (right) along the center.

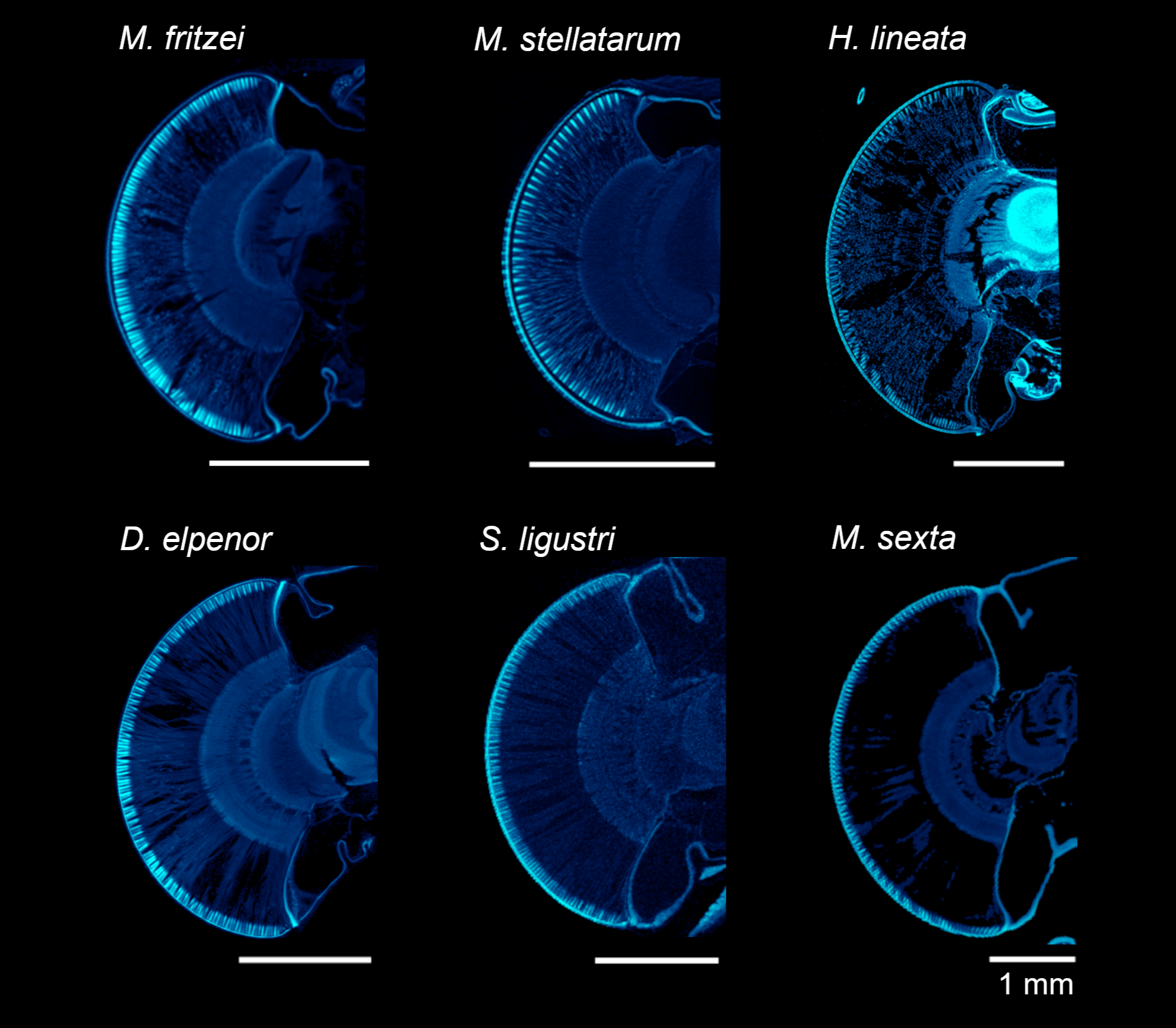

**Figure S6:** Sagittal µCT scan cross-sections of the compound eye of the six hawkmoth species studied. There is no skewness visible when examined in dorsal ventral plane. Each cross-section is in the sagittal plane and crosses through the center of the eye. In all images, the top corresponds to the dorsal side of the head, and the bottom to the ventral.

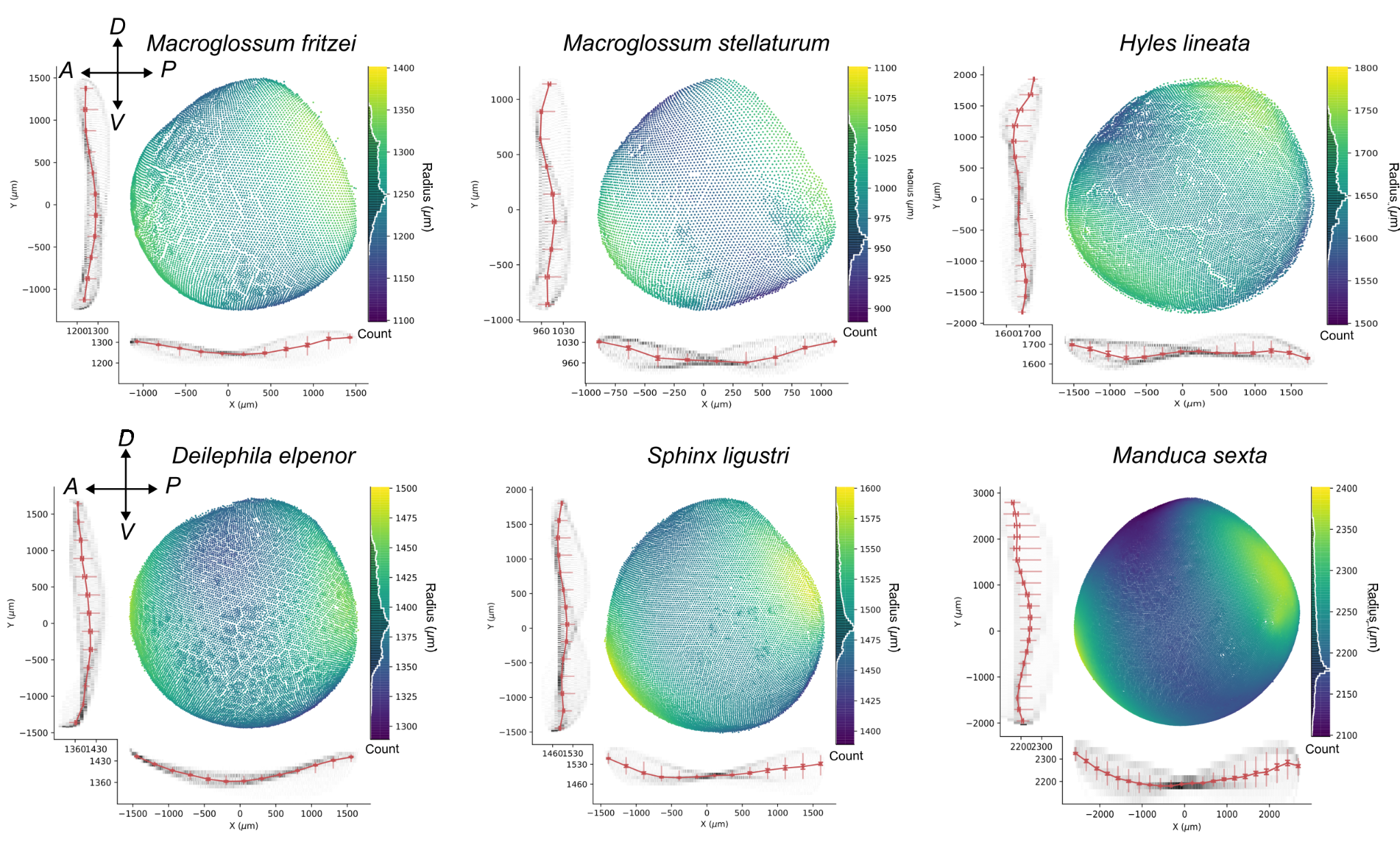

**Fig S7:** Variation in estimated radius from center of eye, in X (ventral to dorsal) and Y (posterior to anterior) for all the hawkmoth species studied. The color heat map ranges are not uniform across all plots but 200-300 μm range around the median was chosen to try and represent somewhat equal variation. Data was rotated to a global coordinate orientation. The red lines represent the number of ommatidia in each bin along these axes.

**Supplememntary Tables**

Supplemental Table 1. Taxon table, status of scan. [Table_S1: CT_eye_list](https://docs.google.com/spreadsheets/d/1_KW3SgW3bXMZwymOdWXsE2V_ElOQ-uHTOb-sef6NRSY/edit#gid=0)

Excel File is uploaded on Figshare

**Table S2**: Manual measurements of Δφ and facet diameter along central to peripheral transects (an average of along the equator -120 degrees and +120 degrees) for the different species, see methods for more detail.

|  | Interommatidial Angle (deg) | | | | | Ommatidia Diameter (µm) | | | | |
| --- | --- | --- | --- | --- | --- | --- | --- | --- | --- | --- |
| Angle (center to peripheral) | 0 | 15 | 30 | 45 | 60 | 0 | 15 | 30 | 45 | 60 |
| *Manduca sexta* | 0.89 | 0.90 | 0.97 | 1.05 | 1.17 | 34.02 | 34.04 | 32.94 | 31.54 | 29.22 |
| *Deilephila elpenor* | 1.35 | 1.35 | 1.41 | 1.49 | 1.57 | 30.08 | 30.16 | 29.46 | 28.18 | 27.44 |
| *Sphinx ligustri* | 1.22 | 1.21 | 1.29 | 1.36 | 1.47 | 30.60 | 30.93 | 29.78 | 28.53 | 28.49 |
| *Hyles lineata* | 1.40 | 1.62 | 1.87 | 1.97 | 2.03 | 31.52 | 32.17 | 32.84 | 33.23 | 33.85 |
| *Macroglossum fritzei* | 1.72 | 1.74 | 1.9 | 2.05 | 2.21 | 31.60 | 31.38 | 30.68 | 30.86 | 30.49 |
| *Macroglossum stellatarum* | 1.85 | 1.95 | 2.07 | 2.19 | 2.25 | 30.02 | 29.58 | 29.67 | 30.04 | 27.89 |

**Table S3**: OLS models of Lens Area as a function of Skewness Angle

| species | F | df | R^2^ | correlation | P | slope | intercept |
| --- | --- | --- | --- | --- | --- | --- | --- |
| *D. elpenor* | 441.538 | 13064.000 | 0.033 | 0.181 | 0.000 | 4.135 | 602.872 |
| *H. lineata* | 684.843 | 11910.000 | 0.054 | 0.233 | 0.000 | 4.091 | 782.728 |
| *M. fritzei* | 975.311 | 8323.000 | 0.105 | 0.324 | 0.000 | 4.813 | 640.290 |
| *M. stellatarum* | 266.015 | 5538.000 | 0.046 | 0.214 | 0.000 | 4.223 | 494.324 |
| *M. sexta* | 0.118 | 28791.000 | 0.000 | 0.002 | 0.731 | 0.026 | 821.411 |
| *S. ligustri* | 0.023 | 12498.000 | 0.000 | 0.001 | 0.881 | 0.022 | 652.233 |

**Supplementary Methods**

**Sampling design and choice of species :**

The hummingbird hawkmoth (*Macroglossum stellatarum)* is a small well-studied hawkmoth that readily feeds in the lab and has been used extensively for foraging experiments (Kelber, Balkenius, and Warrant 2003; Balkenius and Kelber 2004; Telles et al. 2014). *Macroglossum fritzei* exhibits flexible diel activity (Pittaway and Kitching n.d.), but little is known about its life history and ecology. The white-lined sphinx (*Hyles lineata)*, is also active both during the day and night (Broadhead et al. 2017; Aldridge and Campbell 2007) and readily exhibits flower feeding and tethered flight behavior in a lab setting (Kelber, Balkenius, and Warrant 2003; Windsor, Bomphrey, and Taylor 2014). The nocturnal large elephant hawkmoth (*Deilephila elpenor)* is in the same subfamily as hummingbird hawkmoth *Macroglossum*, but more strictly nocturnal, with a greater reliance on odors than visual cues in lab foraging experiments (Hamdorf and Hoglund 1981; Theobald, Warrant, and O’Carroll 2010; Balkenius, Rosén, and Kelber 2006; Johnsen et al. 2006). The tobacco hornworm *(Manduca sexta)* is a large and bulky hawkmoth, with an extremely elongated proboscis. While it has been reported as crepuscular (dusk or dawn active), actual measurements of activity show that can fly all night, so calling it nocturnal is justifiable (Kuenzinger et al. 2019; Broadhead et al. 2017). It is a well-established model in neuroscience, physiology and flight control (Copley, Parthasarathy, and Willis 2018; Stöckl, O’Carroll, and Warrant 2017; Theobald, Warrant, and O’Carroll 2010; Riffell et al. 2013; Putney, Conn, and Sponberg 2019; Treidel et al. 2024; Raguso, LeClere, and Schlumpberger 2005). The other exclusively nocturnal species we used is the privet hawkmoth (*Sphinx ligustri),* an emerging model in behavior and physiology, and although older work has characterized its visual responses neurally, it has more recently been used in flight to light and behavioral experiments (Degen et al. 2024; Storms et al. 2022; Collett 1972).

**Staining procedures and drying**

UF and FLMNH and NHM Staining Protocol used by Yash Sondhi and Debroah Glass
**Removing head and storage**

1. Remove head from moth

2. Remove antennae under light/ dissection microscope

3. Once antennae removed, place head in boiling water for 30 seconds. This helps to prevent brain shrinkage when the head is stored in ethanol for a prolonged period of time. (Not done when only looking at visual structures)

4. Place head in 70% ethanol until the time you wish to stain the head.

Dissection and staining

1. Under a light/dissection microscope remove any excess tissue where the head was removed from the body and remove mouthparts (removing antennae, mouthparts and excess tissue allows the stain multiple entries into the brain).

2. Remove excess ethanol from head with blue roll

3. Place tissue to be stained in a vial with the 1% PTA solution in 70 %ethanol (See recipe below)

4. Store at room temperature for 7 days

5. After 7 days check level of staining using micro-ct

6. If not sufficiently stained place head back into PTA, checking with the micro-ct at regular intervals (e.g. 2 days) to ensure effective staining.

7. Once stained, remove excess PTA solution with blue roll and place back into 70% ethanol until you are ready to scan.

The benefit of using PTA is that it will not overstain the material and it is permanent, meaning the head can be stored once stained. This is unlike for example iodine staining, where a specific length of time in the stain is required to prevent oversaturation and the head must be scanned as soon as possible to prevent the stain leaching out of the head.

PTA is the most effective staining method for visualising the different structures of the brain, with other stains tested the structures were less well defined and thus, more difficult to discern and segment.

**Important**: When scanning, make sure the head is sufficiently blocked to prevent movement during the scan time, otherwise you will have blurry images.

**Note**: If batch scanning to reduce cost, it is advisable to leave samples soaked in water while scanning, to prevent drying.

On average, 1mm of tissue takes 1 days of staining, the larger the sample, the more staining time is required.

Making a 1% PTA solution in 70% ethanol:

1. Make a 1% weight/volume solution of Phosphotungstic acid in water. For creating a 100ml stock solution, dissolve 1 gm of PTA crystals in 100 ml of water.

2. For staining samples, mix 30% of Phosophotungstic acid stock solution with 70% absolute ethanol. For creating a 100 ml stock solution, mix 30 ml of 1% weight/ volume solution with 70 ml of absolute ethanol.

**HDMS protocol Ruchao Quian**
